# Bayesian analysis of resting-state temporal low-γ power in children with phonemic decoding deficits and certain comorbidities

**DOI:** 10.1101/2023.09.20.558564

**Authors:** Oliver Lasnick, Roeland Hancock, Fumiko Hoeft

**Affiliations:** Department of Psychological Sciences, University of Connecticut, Storrs, Connecticut, United States of America; Wu Tsai Institute, Yale University, New Haven, Connecticut, United States of America

**Author notes:** Corresponding author: (OHML).

## Abstract

Dyslexia (decoding-based reading disorder, or RD) is a common learning disorder affecting a large proportion of the population. One theory of the origins of reading deficits is a language network which cannot effectively ‘entrain’ to speech, with cascading effects on the development of phonological skills. Low-gamma (low-γ, 30-45 Hz) neural activity is thought to correspond to tracking at phonemic rates in speech. The main goals of the current study were to investigate temporal low-γ band-power during rest in a sample of children and adolescents with and without RD. We used a Bayesian statistical approach, which has become increasingly popular in the field for its ability to quantify the relative likelihood of competing hypotheses. We examined whether (1) resting-state temporal low-γ power was attenuated in the left temporal region in those with RD; (2) low-γ power covaried with individual performance in reading skills; (3) low-γ temporal lateralization was atypical in the group with RD. Results did not support the hypothesized effects of RD status and poor phonemic decoding ability on left hemisphere low-γ power or lateralization: post-hoc tests revealed that the lack of atypicality in the RD group was not due to the inclusion of those with comorbid attentional deficits. However, post-hoc tests also revealed a specific left-dominance for low-γ rhythms in the RDs only when participants with attentional deficits were excluded. We also observed an inverse relationship between phonemic decoding and left-lateralization in the controls, such that those with better decoding skills were less likely to show left-lateralization, contrary to our expectations. We discuss these unexpected findings in the context of prior theoretical frameworks on temporal sampling, and suggest that our null findings in resting-state EEG may reflect the importance of real-time language processing to evoke gamma rhythms in the phonemic range during childhood and adolescence.

## Introduction

Developmental dyslexia (hereafter referred to as decoding-based reading disorder, or RD) is one of the most common and highly studied specific learning disabilities. Its prevalence in the general population is estimated to be 5-10% [1–2]. Individuals with RD have profound difficulty with reading despite often having normal-range intelligence (IQ) and a lack of other explanatory variables, such as impaired vision [3–4]. RD is highly comorbid with other neurodevelopmental disorders [5–7]. The etiology of RD is thought to be neurobiological, but its specific underlying causes are largely unknown. Cognitive profiles for RD often include deficits in letter-sound knowledge, rapid automatized naming (RAN), and phonological awareness (PA), with PA being the strongest predictor of reading outcomes for English-speaking populations [1, 8]. PA is the ability to explicitly engage one’s knowledge of the component speech sounds (phonemes) which make up a word [9].

One biological perspective suggests that low-gamma (low-γ) activity in the 30-45 Hz range is an important indicator of linguistic processing which may index speech encoding at the phonemic level. Phonemic processing has been shown to be associated with entrainment in the low-γ range during perceptual parsing of speech [10]. Low-γ entrainment may therefore be important for the establishment of strong PA skills: gamma phase-locking has been found to be modulated by phonological contrast in a stimulus, and to show a differential response in typical and poor adolescent readers during sentence-listening [11].

A number of studies have also reported that lateralized intrinsic activity may be related to effective speech processing, phonological processing, and to RD. Resting-state studies in adults have shown that lateralization of low-range gamma band-power in the superior temporal cortices (left hemisphere [LH] > right hemisphere [RH]) is associated with functional asymmetries in speech processing [12]. Intrinsic activity has been shown to predict lateralized language network activity in primary auditory regions [13]. Other studies have reported atypical functional lateralization in individuals with RD, often with weaker relative LH dominance for low-γ activity [14–16]. A recent review proposed that phonemic processing could also be affected by the right frontoparietal attention network, which in turn exerts downstream effects on the LH dorsal reading network [17]. Lateralized low-γ oscillations during resting-state have also been associated with the ability to perceive speech in noise in young children [18].

However, the observed low-γ range in the left superior temporal cortex and its left-right asymmetry have primarily been investigated in children and adults within the context of tasks, such as sentence-listening [11] and auditory steady-state response paradigms [14]. Whether similar trends exist in resting-state EEG is still unknown. This is an important question because endogenous low-γ power, if associated with phonological processing, reading, and RD, would indicate both a potential biomarker and a more generalized processing deficit in the intrinsic auditory and language network in RD. Intrinsic lateralization of resting-state EEG power in the low-γ range in the superior temporal cortices have been reported. In [13], the authors did report that for their sample of adults there is (1) a correlation between fMRI activity and neural oscillations during resting-state at syllabic and phonemic rates; and (2) resting-state lateralization for low-γ activity in the primary auditory cortex; but the latter effect did not extend to the posterior superior temporal cortex. These effects were not found in another study on adults, which reported only a non-significant trend for left-lateralization in the 28-40 Hz range at rest [12].

The literature in children on low-γ band-power during rest is even sparser as it relates to phonological processing, reading, and RD. Gamma activity changes with development, with induced (non-stimulus-locked) amplitudes at 20 and 40 Hz decreasing from age 8 to late adolescence (∼19 years); synaptic pruning, which occurs throughout early development, may be a driving force behind these gamma changes as it decreases unnecessary excitatory-inhibitory dynamics in particular networks [19]. Most relevant, Thompson et al. [18] found left-lateralization of resting low-γ rhythms in children aged 3-5 years that covaried with speech-in-noise perception skills. They suggested endogenous (resting-state) hemispheric specialization for oscillations may support speech processing in challenging listening environments during early childhood. Two other papers also reported links between resting frontal low-γ power and early cognitive and language skills (including nonword reading) in young pre-readers [20–21]. These three studies in children [18, 20–21] sampled pre-readers aged 3-5 and did not look at effects in the superior temporal cortices. These studies also did not examine relationships with phonological processing, reading and (risk for) RD. There is one study to our knowledge that examined resting-state EEG at the low-γ band in RD children: the authors compared cortical sources of eyes-closed resting state EEG at various frequency bands between a small sample of typical readers (TRs) and children with RD [22]. Interestingly, they reported null findings for all bands (including delta, theta, and gamma) except alpha rhythms. Earlier resting-state studies in RD children only examined group differences in the theta and alpha rhythms [23]. Therefore, a more conclusive investigation as to whether there is reduced band-power and atypical lateralization in the temporal lobes that is related to phonological processing, reading, and RD is warranted.

Overall, prior studies on children have shown that there are often PA deficits in RD; that abnormal low-γ phase-locking has been linked to deficits in phonological processing [11]; that those with RD show altered lateralization patterns [15–16, 24]; that reduced left-lateralized entrainment in the low-γ range is correlated with poorer phonological processing and rapid naming in RD [14]; and that there are developmental effects on low-γ power during typical development [19]. These studies are all either behavioral (linking PA to RD) or use task-based designs. What is lacking in the literature is a clear consensus on resting-state dynamics in the reading and language network. Investigating these issues is important because it will inform us about the intrinsic activation of the reading/language network during early to mid-childhood, which may have implications for better understanding children’s language and reading processes.

The aim of the current study is to investigate low-γ power and lateralization at rest in children with and without RD in an age range where children are learning to read (6 to 13 years), expanding prior resting-state studies in beginning readers and adolescents. Analyses were designed to demonstrate whether temporal gamma power is attenuated within the LH in the RD population, reflects reduced left-lateralization compared to TRs, and is predicted by phonemic decoding scores. If gamma power is attenuated in the LH temporal region of RDs, and/or the typical left-lateralization pattern is reduced/reversed compared to TRs, this would suggest that there is inherent functional organization to the brain which localizes processing of hierarchically-structured timescales for auditory stimuli to a dominant hemisphere; and that this hierarchical processing is atypical at phonemic timescales in children with decoding deficits. Additionally, we aim to determine whether any such differences are associated with individual performance in reading-related skills, specifically pseudoword decoding. We predict a positive relationship between low-γ band-power and individual differences in phonemic decoding; and that the effect will be stronger in the left hemisphere compared to the right (LH > RH). This is based on prior work in adults which showed that the LH is more tuned for the processing of gamma band information >20 Hz, while slower delta and theta bands dominate in the right (for a review of oscillation and lateralization differences in RD, see [17]).

## Materials and Methods

### Participants

All data came from the Child Mind Institute Healthy Brain Network Project (CMI HBN), an online repository of multi-site imaging and behavioral data [25]. Neuroimaging and behavioral data from the CMI HBN project are publicly available upon completion of a Data Usage Agreement. They are provided in a de-identified/anonymized format such that none of the authors have access to any information which could be used to re-identify the participants. Therefore, additional participant consent or involvement was not required and no further ethical approval was sought from the University of Connecticut’s Institutional Review Board. Data were accessed for research purposes in May 2020.

The CMI dataset participants range in age from 5-22 years old, however the age range for this study was restricted to those aged 6-13 years. Reasoning was threefold: (1) age 6 is when most children achieve early reading skills such that both reading metrics and individual differences are stable and reliable (standardized scores on tests of word reading may be derived starting at 6, and a majority of children with reading impairments are identified during this time frame), (2) within this age range, PA contributes to individual differences in word-reading and word-reading is reflective of decoding ability [8, 26], and (3) developmental changes in the brain possibly related to neural oscillations in the gamma range have been documented during this time frame [19].

Other inclusion criteria were a full-scale IQ (Wechsler Intelligence Scale for Children [WISC]) score >70 and completion of TOWRE-2 (phonemic decoding [PDE] and sight-word efficiency [SWE] subtests) [27]. TOWRE is a timed word reading measure wherein the participant is presented with a list of either words (SWE) or pseudowords (PDE) and instructed to read each aloud, in order, as quickly and accurately as possible. They are given 45 seconds to do so. Higher scores reflect a larger number of words read and/or pronounced correctly. These two subtests create an overall reading efficiency composite score, which is frequently used as RD classification criteria in research; total scores that fall below 85 (with the expected population average being 100 with ±15 equivalent to 1 SD) are often used to indicate poor reading (for recent examples, see [28–30]).

For the group-based analyses, two main groups were generated: (1) an RD group with both PDE and SWE scores <85, to ensure that the participants had both phonemic decoding and (non-compensated) word-reading deficits; and (2) a TR group who (a) had not received a prior clinician diagnosis of RD and (b) had both TOWRE subtest scores >90. Because the practice of excluding comorbidities or placing restrictions on participant samples (e.g., only including right-handed males) inherently reduces the generalizability of findings - especially in relation to RD, for which comorbidities are highly prevalent and may affect developmental outcomes [31] - we allowed our sample (both RDs and TRs) to have certain clinical diagnoses that often co-occur with reading deficits at elevated rates compared to the general population. We expected that rates of all included diagnoses would be higher in those with RD compared to the TRs. These included depression, anxiety, attention-deficit hyperactivity disorder (ADHD), and other specific learning disorders such as dyscalculia, dysgraphia, and language disorder. Identification of these comorbidities was based on consensus diagnoses from clinicians, following the procedure used by CMI for the HBN project.

This gave a sample size of N = 315 (TRs: N = 192; RD-full: N = 123) before all final processing and quality control/exclusion criteria had been applied (see section on EEG Data Collection and Quality Control below). The frequencies of comorbid diagnoses for all groups are given in Table 1. The statistics in Table 1 are derived from a subsample of N = 261, consisting of those who met both quality control criteria and outlier requirements for the temporal lateralization analyses (see Analyses for more details). Chi-squared tests were performed to determine if the rates of diagnosis for the listed comorbidities differed significantly between the groups: TR vs. RD-full. Because we include common comorbidities, we performed exploratory/secondary analyses comparing RDs with and without the most common comorbidity, ADHD (accounting for 35.0% of the RD-full sample prior to quality control).

**Table 1.** Frequencies of Comorbid Diagnoses (N = 261).

|  | TR | RD-full | X<sup>2</sup> (p) | RD-only | RD-ADHD | X<sup>2</sup> (p) |
| --- | --- | --- | --- | --- | --- | --- |
| *ADHD Subtypes* |  |  |  |  |  |  |
| ADHD-Combined Type | 0 | 18 | **31.13 (<.001)** | 0 | 18 | **40.77 (<.001)** |
| ADHD-Inattentive Type | 0 | 16 | **27.44 (<.001)** | 0 | 16 | **35.37 (<.001)** |
| ADHD-Hyperactive/Impulsive Type | 0 | 1 | (.383)<sup>a</sup> | 0 | 1 | (.350)<sup>a</sup> |
| *Specific Learning Disorders* |  |  |  |  |  |  |
| Reading | 0 | 37 | **69.41 (<.001)** | 23 | 14 | .208 (.648) |
| Mathematics | 3 | 3 | (.678)<sup>a</sup> | 2 | 1 | (1.00)<sup>a</sup> |
| Written Expression | 1 | 3 | (.159)<sup>a</sup> | 2 | 1 | (1.00)<sup>a</sup> |
| Language Disorder | 8 | 14 | **6.52 (.011)** | 8 | 6 | (.553)<sup>a</sup> |
| Speech Sound Disorder | 3 | 0 | (.288)<sup>a</sup> | 0 | 0 |  |
| *Anxiety Disorders* |  |  |  |  |  |  |
| Generalized Anxiety Disorder | 15 | 2 | **5.42 (.020)** | 2 | 0 | (.540)<sup>a</sup> |
| Specific Phobia | 8 | 4 | (1.00)<sup>a</sup> | 4 | 0 | (.295)<sup>a</sup> |
| Separation Anxiety | 7 | 3 | (.746)<sup>a</sup> | 3 | 0 | (.550)<sup>a</sup> |
| Social Anxiety (Social Phobia) | 8 | 1 | (.160)<sup>a</sup> | 0 | 1 | (.350)<sup>a</sup> |
| Other Specified Anxiety Disorder | 10 | 3 | (.381)<sup>a</sup> | 2 | 1 | (1.00)<sup>a</sup> |
| *Depressive Disorders* |  |  |  |  |  |  |
| Persistent Depressive Disorder (Dysthymia) | 1 | 0 | (1.00)<sup>a</sup> | 0 | 0 |  |
| Major Depressive Disorder | 2 | 0 | (.525)<sup>a</sup> | 0 | 0 |  |
| Other Specified Depressive Disorder | 1 | 0 | (1.00)<sup>a</sup> | 0 | 0 |  |
| *Conduct Disorders* |  |  |  |  |  |  |
| Oppositional Defiant Disorder | 0 | 0 |  | 0 | 0 |  |
| Intermittent Explosive Disorder | 0 | 0 |  | 0 | 0 |  |
| Other Specified Disruptive, Impulse-Control, and Conduct Disorder | 0 | 0 |  | 0 | 0 |  |
| *Other Attentional Issues Not Meeting ADHD Criteria* |  |  |  |  |  |  |
| Unspecified Attention-Deficit/Hyperactivity Disorder | 3 | 0 | (.288)<sup>a</sup> | 0 | 0 |  |
| Other Specified Attention-Deficit/Hyperactivity Disorder | 12 | 4 | 1.28 (.258) | 4 | 0 | (.295)<sup>a</sup> |
| Adjustment Disorders | 4 | 1 | (.652)<sup>a</sup> | 0 | 1 | (.350)<sup>a</sup> |

ADHD, attention deficit hyperactivity disorder.

For the TR vs. RD-full comparisons, as expected, differences in diagnostic frequency emerged for ADHD, language disorder, and specific learning disorders of reading, where the RD-full group had significantly greater diagnostic rates than TRs (all ps <.05). Unexpectedly, the reverse effect was observed for Generalized Anxiety Disorder, where TRs were diagnosed at greater rates compared to RDs (p <.05). For the RD-only vs. RD-ADHD comparisons, the RD-ADHD group had a self-evidently higher rate of ADHD diagnoses, while all other comparisons were non-significant (ps >.1).

Missing socioeconomic status (SES) data was imputed using the multiple imputation method as described in [32], in order to control for non-random missingness of data and to preserve statistical power. The multiple imputation method was chosen to preserve population variance, as previous studies have shown that SES is correlated with multiple measures of cognitive functioning, including reading achievement [33–34]. SES values coded for self-reported annual income on an ordinal scale of 0 (< $10,000/year) to 11 (> $150,000/year). Prior to imputation, N = 254 (80.6%) participants had SES data available, meaning that a total of 61/315 SES scores were imputed. Missing SES data was imputed using the variables FSIQ, TOWRE PDE scores, and group (RD vs. TR status) as predictors, as these were significantly correlated with the existing SES data. Table 2 describes demographic and behavioral variables, again for the N = 261 sample for the temporal lateralization analyses.

**Table 2.**
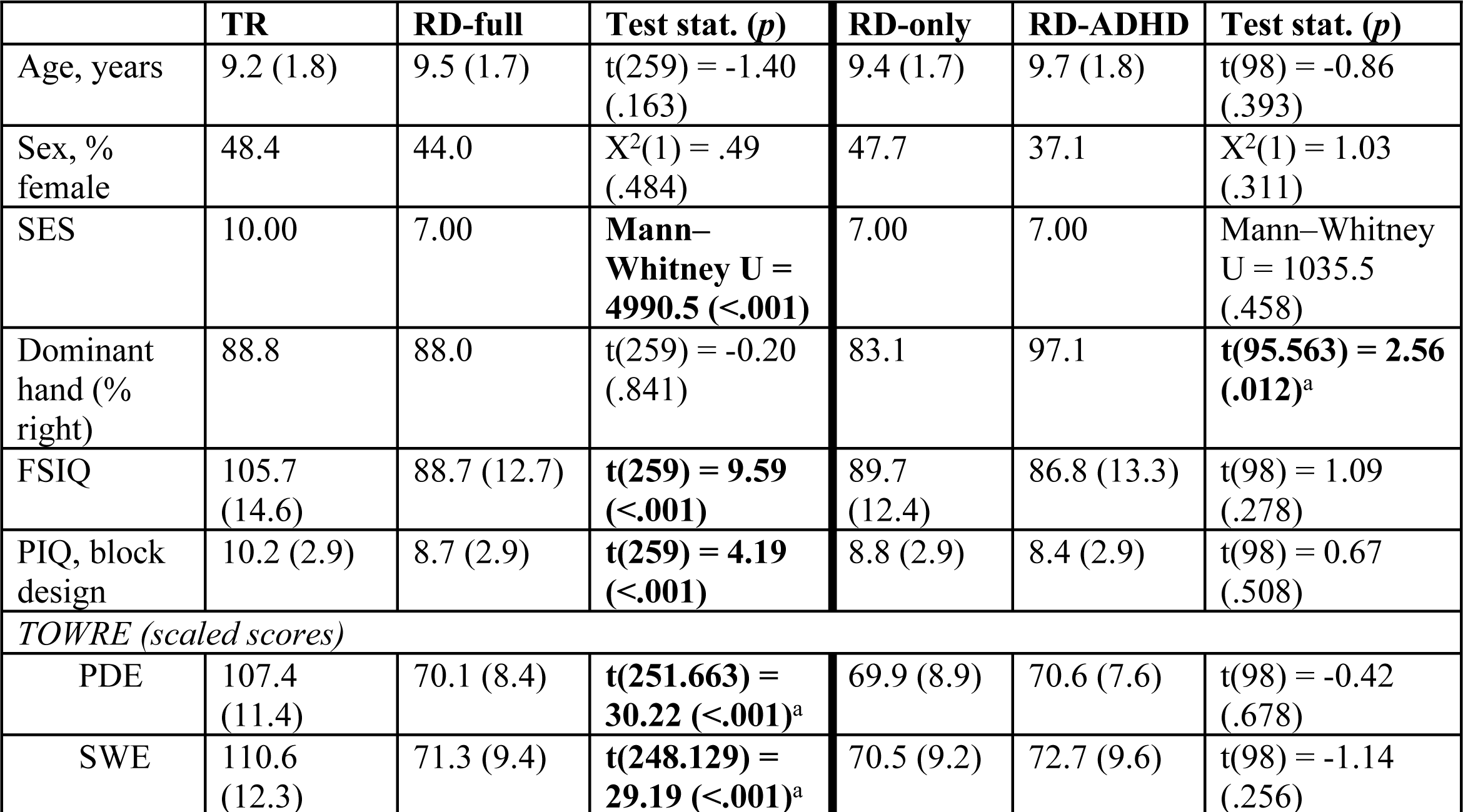
Demographics of sample (N = 261).

|  | TR | RD-full | Test stat. (*p*) | RD-only | RD-ADHD | Test stat. (*p*) |
| --- | --- | --- | --- | --- | --- | --- |
| Age, years | 9.2 (1.8) | 9.5 (1.7) | $t(259) = -1.40$<br>(.163) | 9.4 (1.7) | 9.7 (1.8) | $t(98) = -0.86$<br>(.393) |
| Sex, % female | 48.4 | 44.0 | $X^2(1) = .49$<br>(.484) | 47.7 | 37.1 | $X^2(1) = 1.03$<br>(.311) |
| SES | 10.00 | 7.00 | **Mann–Whitney U = 4990.5 (<.001)** | 7.00 | 7.00 | Mann–Whitney U = 1035.5<br>(.458) |
| Dominant hand (% right) | 88.8 | 88.0 | $t(259) = -0.20$<br>(.841) | 83.1 | 97.1 | **$t(95.563) = 2.56$**<br>**(.012)<sup>a</sup>** |
| FSIQ | 105.7<br>(14.6) | 88.7 (12.7) | **$t(259) = 9.59$**<br>**(<.001)** | 89.7<br>(12.4) | 86.8 (13.3) | $t(98) = 1.09$<br>(.278) |
| PIQ, block design | 10.2 (2.9) | 8.7 (2.9) | **$t(259) = 4.19$**<br>**(<.001)** | 8.8 (2.9) | 8.4 (2.9) | $t(98) = 0.67$<br>(.508) |
| *TOWRE (scaled scores)* |  |  |  |  |  |  |
| PDE | 107.4<br>(11.4) | 70.1 (8.4) | **$t(251.663) = 30.22$**<br>**(<.001)<sup>a</sup>** | 69.9 (8.9) | 70.6 (7.6) | $t(98) = -0.42$<br>(.678) |
| SWE | 110.6<br>(12.3) | 71.3 (9.4) | **$t(248.129) = 29.19$**<br>**(<.001)<sup>a</sup>** | 70.5 (9.2) | 72.7 (9.6) | $t(98) = -1.14$<br>(.256) |

SES, socioeconomic status; FSIQ, full-scale IQ; PIQ, performance IQ; TOWRE, Test of word reading efficiency; PDE, phonemic decoding efficiency; SWE, sight-word efficiency.

### Electrophysiological (EEG) Data Collection

#### Resting-state paradigm

Participants sat in front of a computer and viewed a fixation cross displayed in the center of the screen (Fig. 1). They were given the following instructions: ‘Fixate on the central cross. Open or close your eyes when you hear the request for it. Press to begin.’ During the paradigm they were instructed by a recorded voice from a female research assistant to open or close their eyes at various points throughout the run. A total of 10 segments were collected with eyes open (EO) or closed (EC) at alternating times (5 segments for both the EO and EC conditions). Each EO segment was 20 seconds long; each EC segment was 40 seconds long. Both EO and EC data were used for the analyses (total of 300 seconds). The entire paradigm lasted 5 minutes.

**Fig 1.**
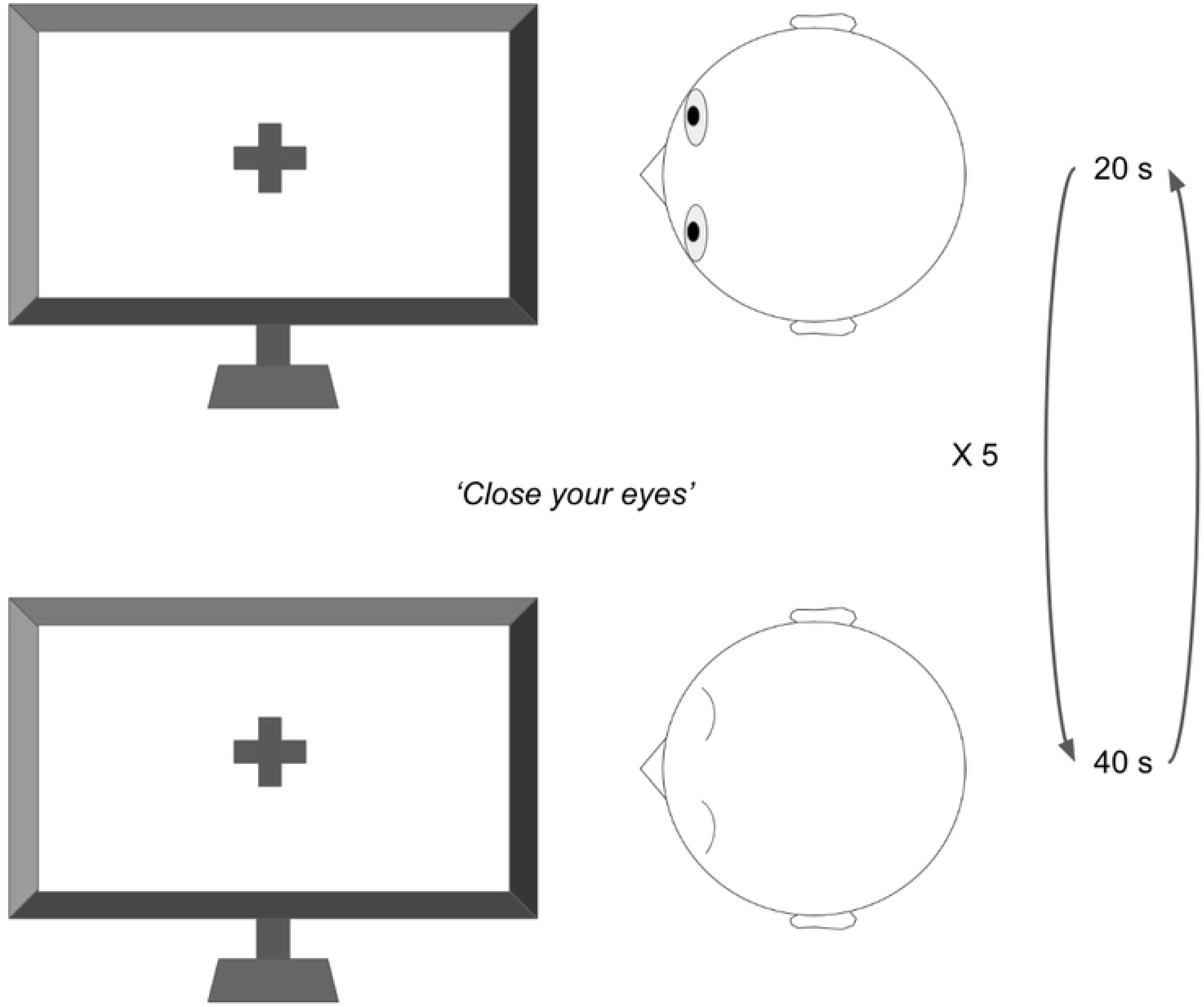
Illustration of resting-state paradigm. Participants are asked to close/open their eyes intermittently for a total of 300 seconds.

#### Acquisition

Data were acquired from a high-density 128-channel net using the EGI (Electrical Geodesics, Inc.) Geodesic Hydrocel system. Collection was performed in a sound-attenuated room with a sampling rate of 500 Hz and on-line 0.1-100 Hz bandpass filter. The recording reference was Cz. Simultaneous eye tracking was performed by recording eye position and pupil dilation with an infrared video-based eye tracker at a sampling rate of 120 Hz. 9 EOG channels were placed on the forehead, outer and inner canthi (electrodes E8, E14, E17, E21, E25, E125, E126, E127, and E128).

#### Preprocessing

All data was preprocessed using Automagic, a MATLAB-based toolbox and EEGLAB plugin for automatic preprocessing and quality assessment of large-scale EEG data [35]. First, individual bad channels were identified and all had line noise removed using the PREP pipeline [36]. An off-line high-pass 1 Hz filter was used. It has been suggested that a high-pass filter of 1-2 Hz produces ideal signal-to-noise ratio when utilizing Independent Component Analysis (ICA), and that this approach often works better than electrooculogram-based ocular artifact regression [37]. An offline low-pass 55 Hz filter was also selected.

#### Quality control (QC)

ICA was done with ICLabel [38]. Artifacts associated with muscle, heart, and eye activity were removed. Individual channels flagged by PREP and channels with excessively high or low variance were discarded. Residual bad channel detection was run after the entire preprocessing pipeline finished to catch any remaining channels with excessive variance. Bad channels were interpolated using the spherical method. After all preprocessing and SES imputation, 338 participants’ files were loaded into MNE-Python [39]. Both EO and EC resting-state data were re-epoched into 1-second epochs and had their last second discarded, for a total of up to 300 1-s epochs per subject. Bad epochs were identified and 100 ‘good’ epochs (based on built-in MNE standards for epoch rejection) used as the minimum for subject inclusion; only good epochs had their gamma power calculated (see next section). Of this sample, 23 subjects had at least one missing TOWRE subtest score and were excluded from further analysis, reducing the sample to N = 315. Finally, the minimum-epoch criteria was applied and outliers were excluded for each dependent variable in their respective analysis.

#### Calculation of low-γ power

For this study temporal electrodes were selected a priori due to their relevance to the reading and language network. In addition, some adult resting-state literature reports that intrinsic hemispheric asymmetries in low-γ power may be localized to auditory cortex and/or superior temporal cortices [12–13]. Electrodes on the outer rim and nasion were excluded. The final sets of electrodes were the following: ‘E34’, ‘E35’, ‘E39’, ‘E40’, ‘E41’, ‘E45’, ‘E46’, ‘E47’, ‘E50’, ‘E51’, and ‘E58’, for the left hemisphere; and ‘E96’, ‘E97’, ‘E98’, ‘E101’, ‘E102’, ‘E103’, ‘E108’, ‘E109’, ‘E110’, ‘E115’, and ‘E116’ for the right hemisphere (Fig. 2).

**Fig 2.**
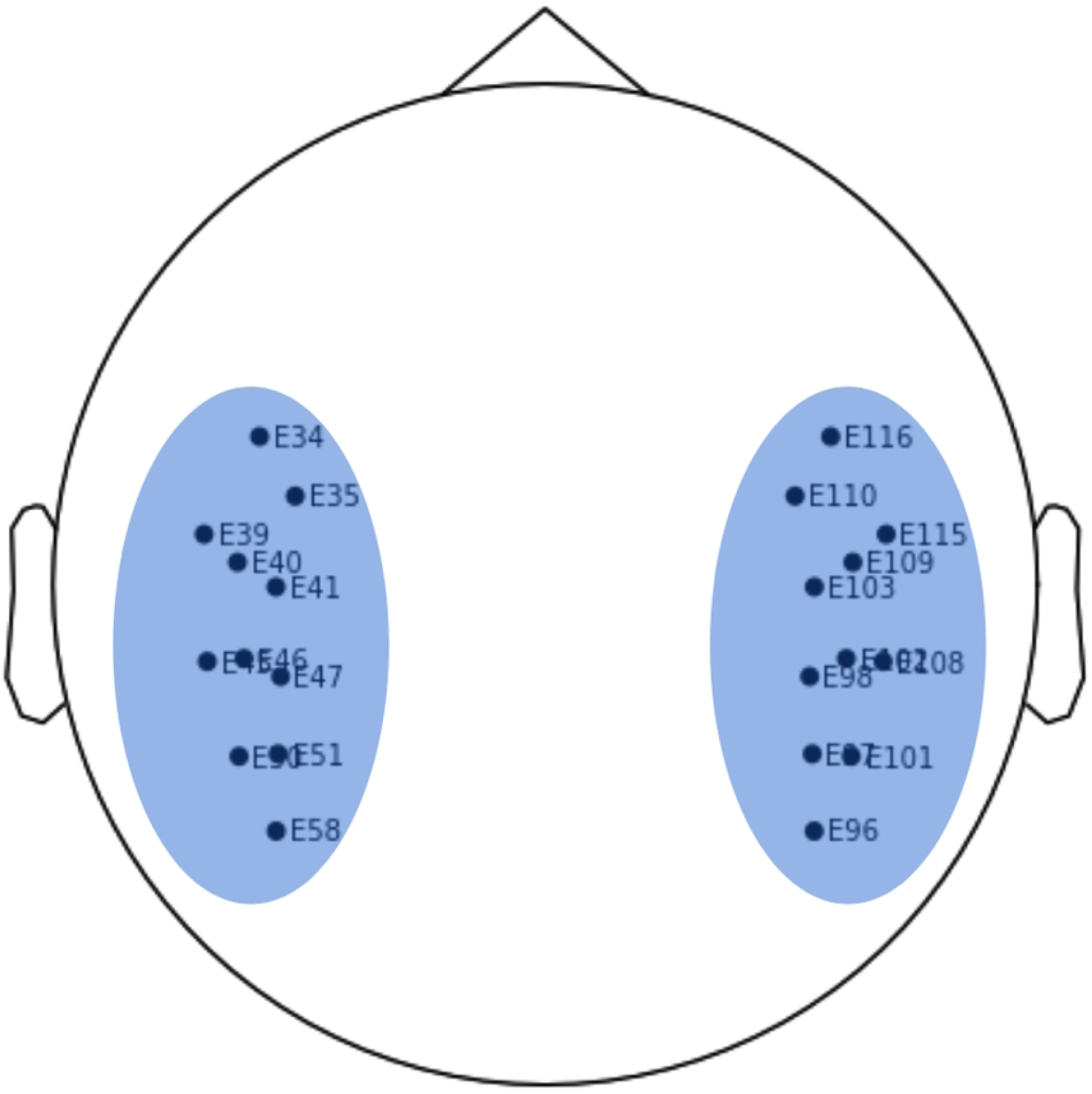
Chosen electrodes for analyses.

For each channel, power spectral density (PSD) was estimated using the multitaper method as implemented in the MNE-Python package. Mean gamma band-power in the 30-45 Hz range was calculated for all epochs, averaged across epochs, and subjected to a log transform of 10 * log_10_(x) to derive absolute power in units of μV^2^/Hz (dB). These values were then averaged for the left and right hemisphere temporal lobes.

### Analyses

#### Outliers

Outlier analysis and exclusion were performed in SPSS Statistics v28. Outliers in SPSS fall outside the range [1^st^ quartile - (1.5 × IQR), 3^rd^ quartile + (1.5 × IQR)], where IQR is the interquartile range. The resulting distributions of each dependent variable were visually examined and judged to be approximately normal. This left a final N = 245 (17 outliers removed) for the temporal analyses and N = 261 (1 outlier removed) for the temporal lateralization analyses. For both repeated measures (RM) ANCOVAs, Box’s Test for equality of covariance matrices and Levene’s Test for equality of error variances were run in SPSS prior to the analyses. Only Levene’s Test was used for the non-RM ANCOVA and regression models. An alpha threshold of .05 was used for all frequentist significance tests (Box’s and Levene’s), and effect size was calculated as η^2^.

#### Bayesian statistics

Bayesian analysis, or Bayes factor analysis, is based on Bayes’ Rule, which presents the following mathematical relationship between the probabilities of two events A and B:

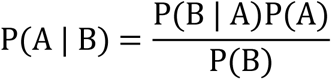

We can therefore state that the probability of a hypothesis H given the observed data is proportional to the probability of observing the data given that H is true, multiplied by the prior probability of H: P(H | data) ∝ P(data | H)P(H).

Bayes factor analysis was conducted in JASP v0.14.1 (jasp-stats.org). The advantage of Bayesian analysis as opposed to frequentist approaches is that it allows researchers to quantify the relative predictive performance of two competing models for the observed data. The Bayes factor (BF_10_) is the ratio between the probabilities of observing the data given that the alternative hypothesis (H_1_) is true, and observing the data given that the null hypothesis (H_0_) is true:

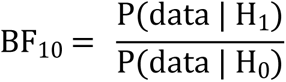

A BF_10_ <.3 indicates evidence favoring the null hypothesis, while a BF_10_ >3 indicates evidence in favor of the alternative hypothesis; values in between indicate that evidence is either anecdotal (.3 < BF_10_ < 3) or absent (BF_10_ ≈ 1). The inclusion Bayes factor (BF_incl_) is also calculated for all individual factors in each model to determine the contribution of individual effects. The BF_incl_, when calculated across matched models, estimates an effect’s unique contribution by comparing all models that contain the effect to equivalent models without it:

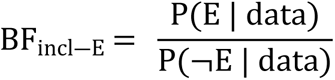

where P(E | data) + P(¬E | data) = 1. Additionally,

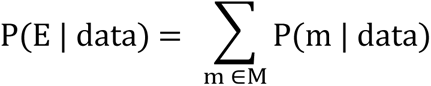

where M is the set of all models *m* containing the effect E.

#### Analysis Plan

For analyses where a comparison between the TR and RD-full groups was of interest, post-hoc Bayesian analyses were conducted for each RD subgroup if there was a meaningful main effect of or interaction involving the factor Group, such that the effect was 2x as likely to be included as excluded (BF > 2).

1.1. To examine group differences in temporal low-γ power between TRs and RDs and their interactions with hemisphere, we use a repeated-measures analysis of covariance (RM-ANCOVA) with Hemisphere as a within-subjects factor and Group (TR, RD-full) as a between-subjects factor. Covariates were age, SES, PIQ, sex, and handedness.
1.2. To examine individual differences in temporal gamma power across decoding skills, associations between TOWRE PDE scores and low-γ power were analyzed with two separate multiple linear regression analyses collapsed across group: TOWRE PDE was the independent variable, while LH and RH temporal low-γ power were dependent. Collinearity diagnostics were run in SPSS prior to the analysis to determine which covariates to retain, in order to ensure there was no excessive collinearity with PDE.
2.1. To examine lateralization of intrinsic low-γ power in the temporal lobe, the lateralization index (LI) for low-γ power was quantified according to the same formula used in [18]: (Left - Right) / (Left + Right). One-tailed one-sample t-tests were run on both groups (TR, RD-full) to determine if low-γ power was left-lateralized in the temporal lobes (LI > 0).
2.2. Hemispheric asymmetries were examined for their relationship to phonemic decoding by regressing the LI on PDE collapsed across groups. Covariates are the same as in 1.2.
3.1. To confirm that there are no significant differences between RD-only and RD-ADHD groups in temporal low-γ power in either hemisphere, we perform an exploratory RM-ANCOVA with Hemisphere as a within-subjects factor and Group as a between-subjects factor (RD-only and RD-ADHD groups are the two levels for the between-subjects factor), and temporal gamma power as the dependent variable. Covariates were age, SES, PIQ, sex, and handedness.

## Results

### 1. Temporal low-γ band-power

#### 1.1. RD-full vs. TR

When analyzing band-power in the RD-full and TR groups, Box’s test was not significant (F(3,1615099.27) = 1.526, Box’s M = 4.622, p =.206). Both LH and RH temporal gamma power passed Levene’s test (LH: F(1,243) = 1.843, p =.176; RH: F(1,243) = .166, p =.684). There was no effect of the covariate age (BF _incl-Age_ = .785) when examining data from both groups. Contrary to the hypotheses, when the covariates were added to the null model, results from the Bayesian RM-ANCOVA did not show a BF_10_ > 1 for any of the alternative models with the effects of interest: not Group (BF_10_ = .461), Hemisphere (.202), nor the model with both of the additive effects plus the interaction of Group x Hemisphere (.017) (see Fig. 3). Inclusion Bayes factors were similarly low: BF_incl-Group_ = .463, BF_incl-Hem_ = .204, and BF_incl-Group x Hem_ = .179. The posterior model probability was also highest for the null model with covariates: P(H_0_ | data) > P(H_i_ | data) for all of the alternative models H_i_.

**Fig 3.**
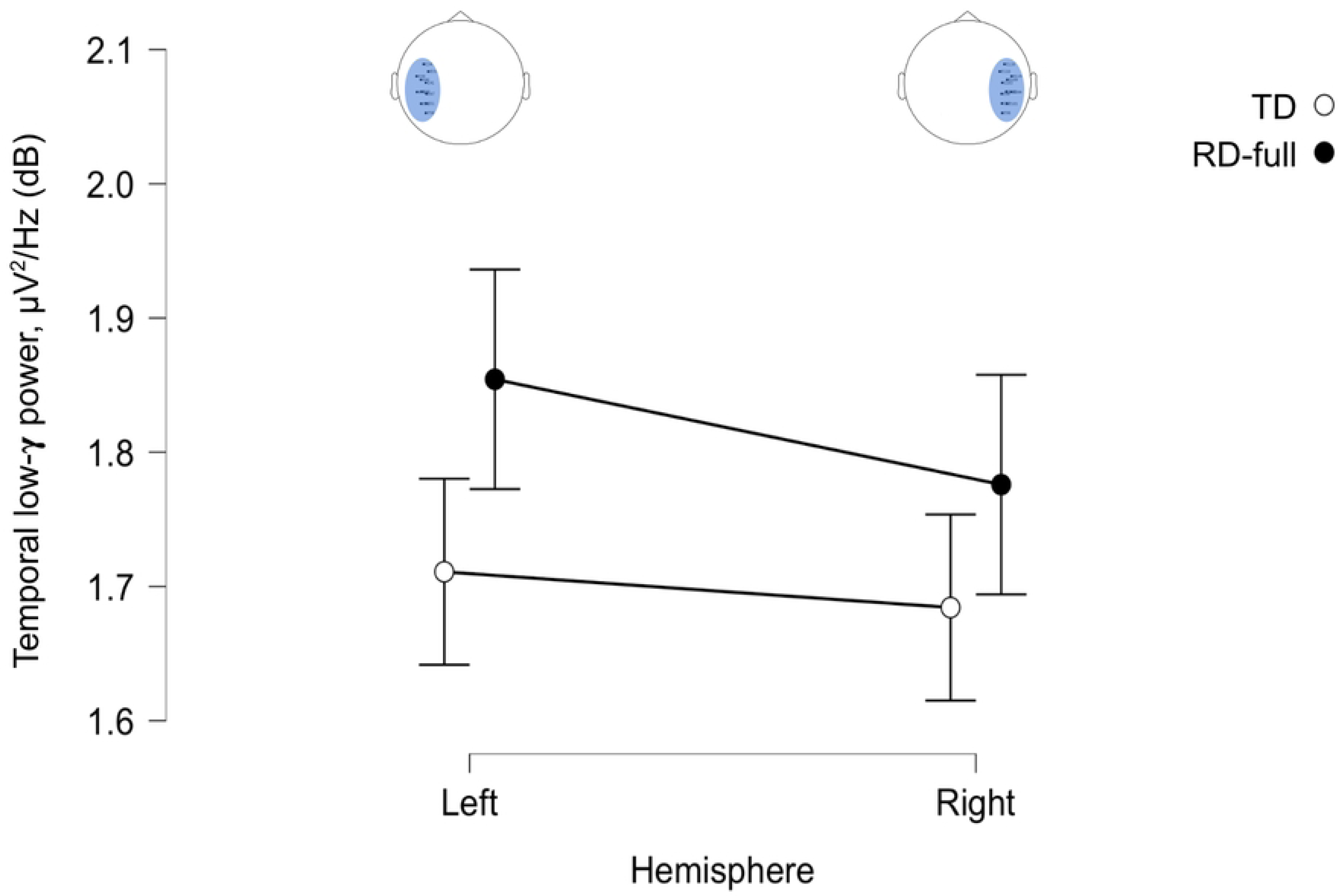
Plots of temporal low-γ power from both hemispheres in the TR and RD-full groups. Error bars are 95% credible intervals.

#### 1.2. Regression of LH/RH low-γ power on phonemic decoding scores

After running collinearity diagnostics, included covariates were sex and handedness: the planned covariates PIQ, SES, and age were excluded due to high collinearity with PDE.

LH: The model with the best relative predictive performance included only Sex, BF_10_ = 6442.424 (females < males). After observing the data, the odds of including the effect of PDE decreased by a factor of 2.865 relative to the prior (BF _incl-PDE_ = .349); its marginal inclusion probability also decreased, suggesting there is likely no unique contribution of PDE to left temporal low-γ power. There was a similar lack of effect for handedness (Fig. 4).

**Fig 4.**
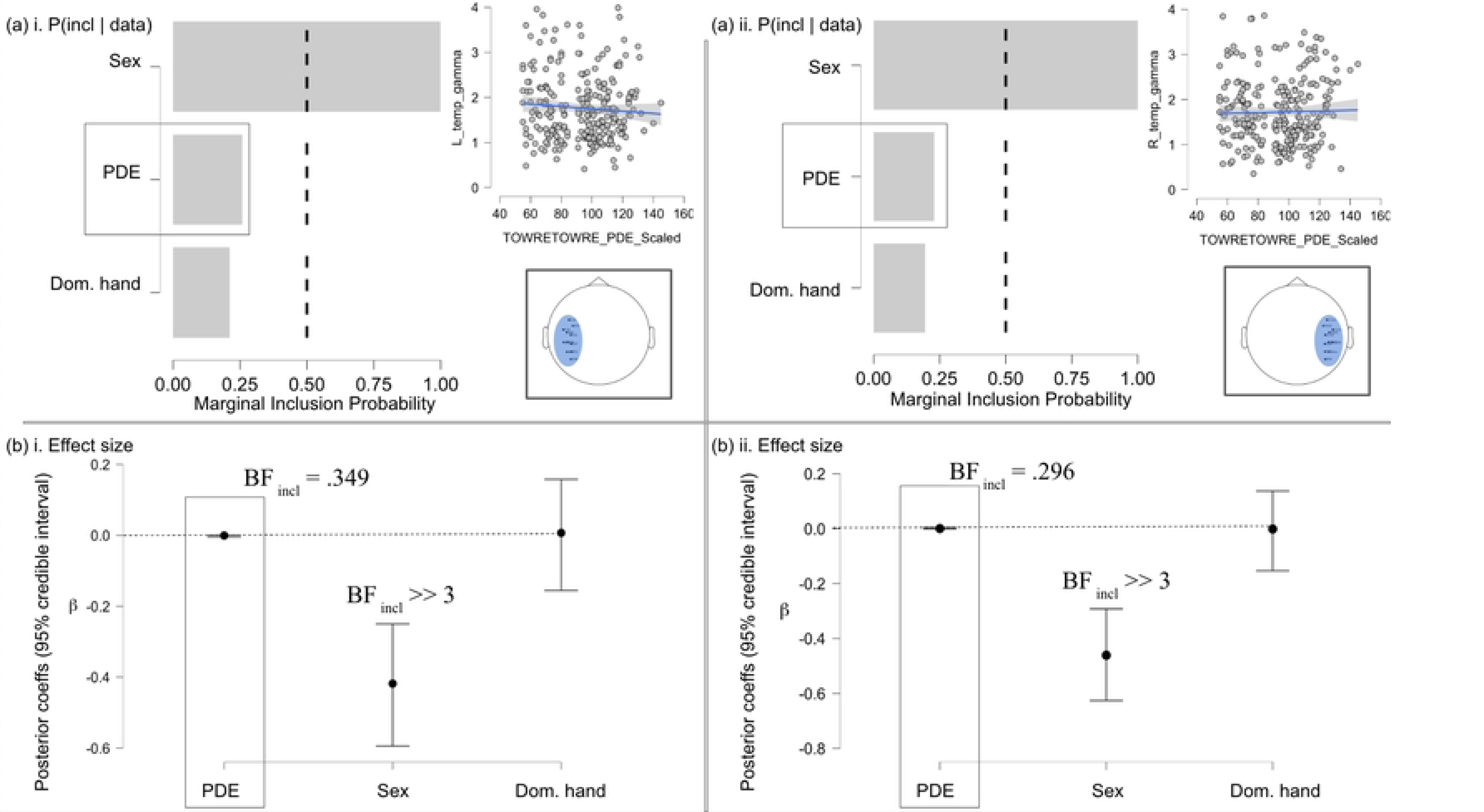
Descriptive plots, marginal inclusion probabilities [P(incl-E | data)], and posterior coefficients (Bayesian measures of effect size). (a) Marginal inclusion probabilities (MIPs) for all effects from separate regression models for the left (a-i) and right (a-ii) hemispheres; MIPs reflect the value of the updated priors (initial values of .50, reflecting a 50-50 chance of inclusion before observing data) for each individual effect after observing the data. (b) Measures of effect size (β coefficients) for the left (b-i) and right (b-ii) hemispheres. The independent variable of interest (PDE) is emphasized despite showing no effect. Error bars are 95% credible intervals.

RH: Once again the most predictive model included Sex (females < males) and no other predictors (BF_10_ = 41403.653), with there likely being no contribution of PDE (or handedness) to right temporal low-γ power: BF _incl-PDE_ = .296.

### 2. Lateralization index

#### 2.1. Lateralization indices in each subgroup

Results indicated slightly different lateralization indices between subgroups. A Bayesian one-sample one-tailed t-test in the positive direction generates values for BF_+0_ and its inverse, BF_0+_, where BF_+0_ = 1 / BF_0+_. Both describe the relative likelihoods of the alternative hypothesis (+, the hypothesis where LI > 0, indicating left-lateralization at rest) and the null (LI = 0, no lateralization at rest).

TRs: A Bayesian one-sample one-tailed t-test resulted in a BF_0+_ = 3.173 for the typical readers. This indicates that the observed data are approximately 3x as likely under the null hypothesis (no lateralization) for the control group.

RD-full: An identical one-sample one-tailed Bayesian t-test generated a BF_0+_ of 1.066 for the RD-full group, suggesting equal relative likelihood of the null and alternative hypotheses (anecdotal evidence).

RD-only (exploratory): We perform a post-hoc follow-up analysis excluding the comorbid RD-ADHD participants due to the ambiguous result reported for the RD-full group. In contrast to the previous findings, results in the RD-only group generated a BF_+0_ = 1.660, suggesting that the data are 1.66x more likely under the alternative hypothesis of left-lateralization than the null.

In summary, given the observed data, the null hypothesis of absent resting-state lateralization for low-γ power was moderately better at predicting the observed data for the TRs than the alternative (left-lateralization). Evidence for the RD-full group was not conclusive for either the null or alternative hypothesis (BF ≈ 1). The subsequent test in the RD-only group suggested, in contrast, that the alternative hypothesis of LI > 0 was comparatively better at explaining the data than the null for this subgroup (Fig. 5). Robustness checks based on changes in the specification of priors, and sequential analysis based on the number of observations (accumulated evidence), are provided in the supplementary material (S1, S2 Figs).

**Fig 5.**
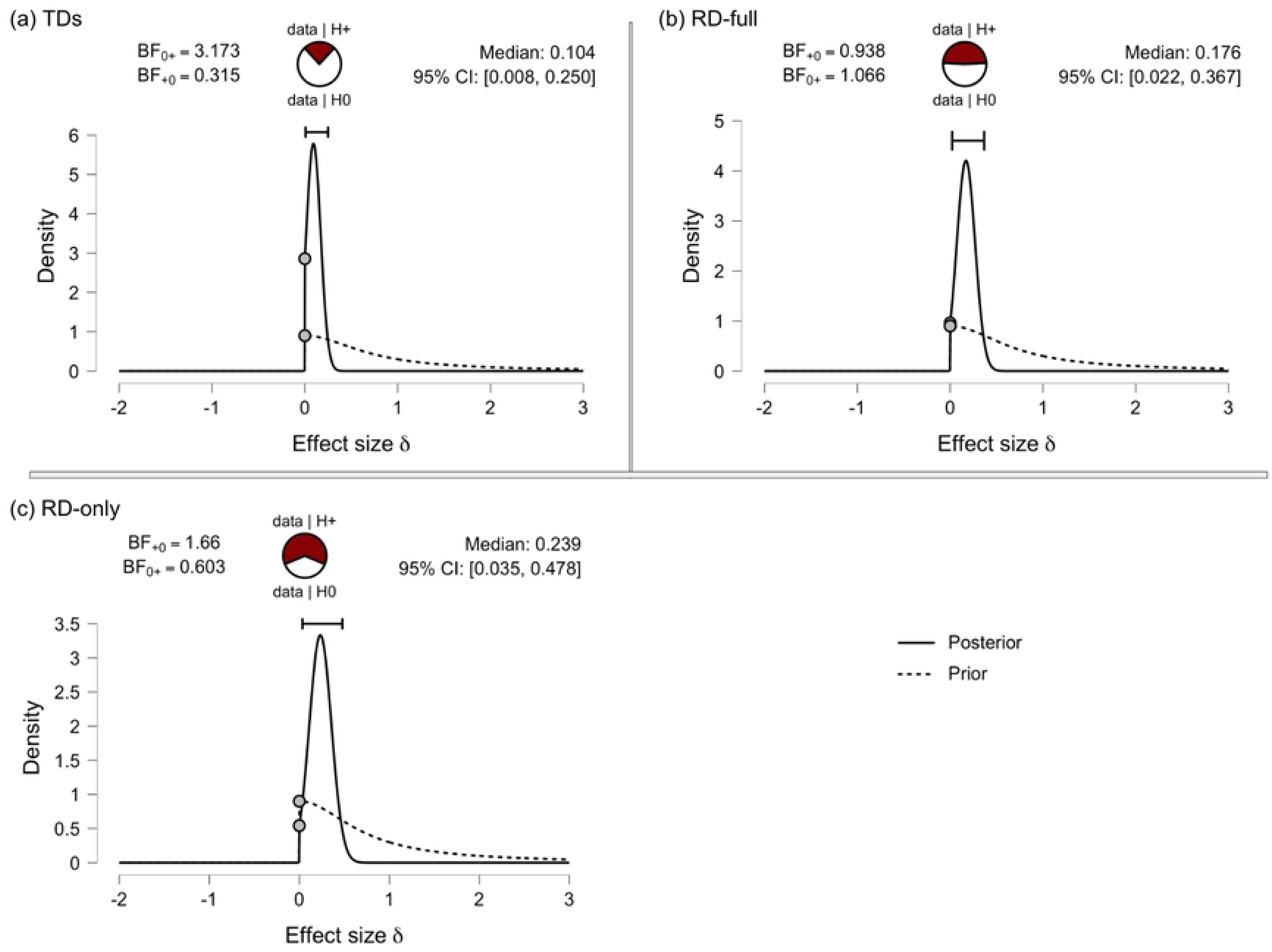
A visual representation of the Bayesian results from the one-sided one-sample t-tests. Each plot shows the Bayes factor (upper left) and relative predictive ability of the null vs. alternative hypothesis (upper center). Density plots show both the prior (before updating beliefs) and posterior (after updating beliefs with observed data) of effect size estimates. CIs are 95% credible intervals. Data comes from (a) TRs; (b): RD-full group; (c) RD-only subgroup.

#### 2.2. Regression of hemispheric asymmetry on PDE

Included covariates were sex and handedness. None of the alternative models had a sufficiently large Bayes factor to support their relative likelihood over the null model (all BF_01_s > 1). The best-performing model relative to the null was the one which included phonemic decoding skill as the sole predictor of LI (BF_10_ = .416, BF_01_ = 2.402); all others had BF_01_s > 3.

*Exploratory post-hoc regression analyses.* The TR and RD-full groups were analyzed separately in an exploratory post-hoc analysis. For the RD-full group all models had BF_01_s > 1. The BF_incl-PDE_ was.330 and P(incl-PDE | data) < .5, suggesting a lack of a strong relationship between PDE and LI for the RD group. For the TR group, the best-performing model had PDE as the sole predictor and had a BF_10_ = 1.631, meaning that the probability of observing the data was slightly higher under this model than the null. For the TRs, but not the RDs, lower PDE scores were associated with left-lateralization (Fig. 6). However, the BF_incl-PDE_ across all possible models was still less than 1 (.823), meaning that P(incl-PDE | data) < P(excl-PDE | data). Given the observed data, the null model was still more likely than the sum of the alternative models that include PDE: P(H_0_ | data) = .484 > P(incl-PDE | data) = .452.

**Fig 6.**
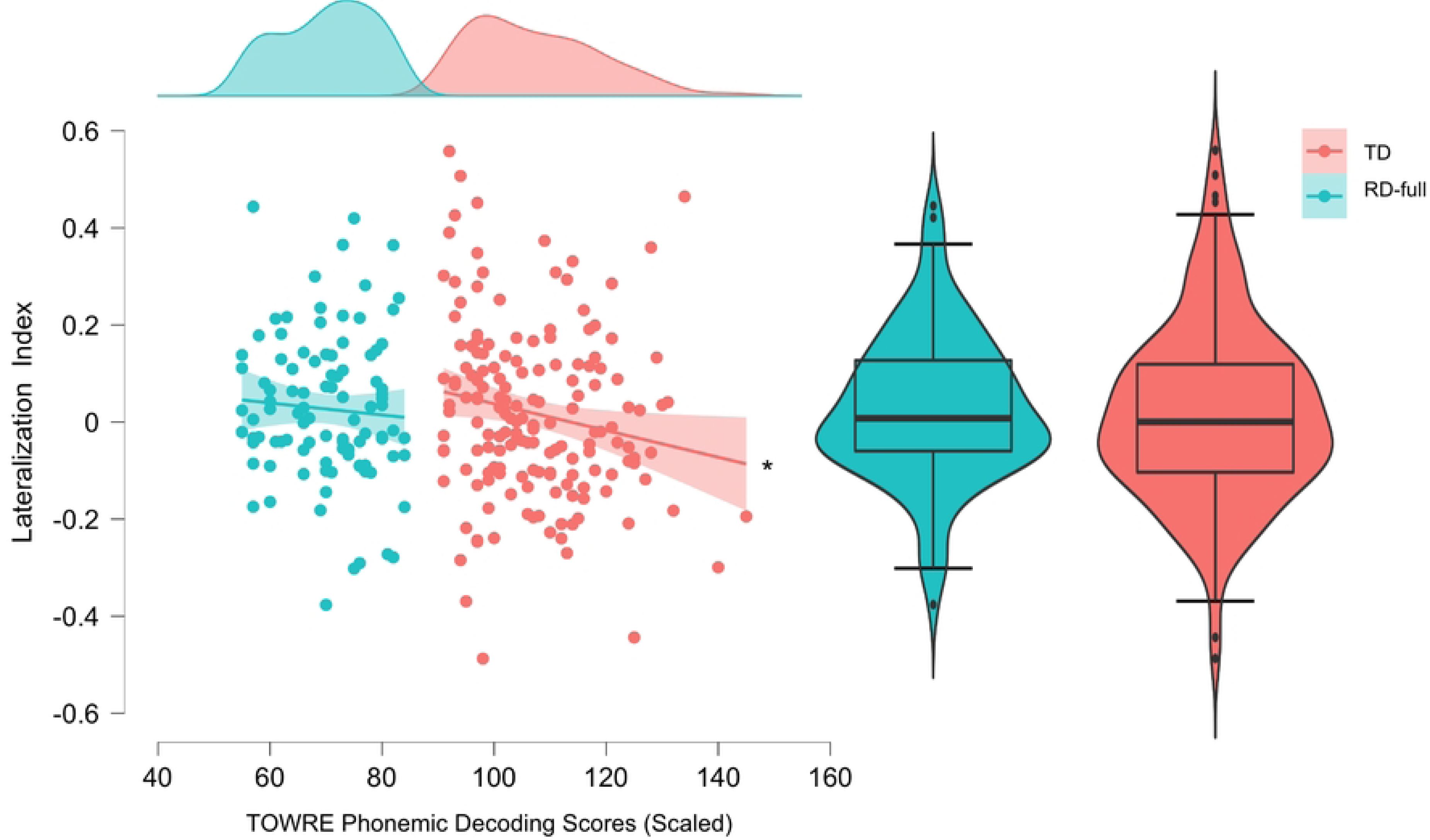
Descriptive plots illustrating the relationship between TOWRE phonemic decoding and lateralization index (LI). A star (*) indicates that the BF_10_ for the model including PDE is >1. Error bars are 95% confidence intervals.

### 3. Exploratory: RD-only vs. RD-ADHD

With regards to band-power in the RD-only and RD-ADHD subgroups, Box’s test for equality of covariance matrices of the dependent variables was not significant for the RM ANCOVA model (F(3,144979.921) = .362, Box’s M = 1.114, p =.781). Both LH and RH temporal gamma power passed Levene’s test for equality of error variances across groups (LH: F(1,92) = .008, p =.928; RH: F(1,92) = .484, p =.488).

For the main Bayesian RM-ANCOVA analysis, the covariates were age, SES, PIQ, sex, and handedness. There was no effect of the covariate age (BF _incl-Age_ = .551) on band-power in the temporal lobes for the RD group. When the covariates were added to the null model, as expected, all BF_01_s > 1.5 for all possible alternative models (Group + intercept, Hemisphere + intercept, Group + Hemisphere + intercept, and Group + Hemisphere + Group*Hemisphere + intercept), meaning that the data was at least 1.5x more likely to be observed under the null compared to any of the alternative hypotheses.

## Discussion

The goal of this study was to analyze differences in resting-state low-γ band-power associated with decoding-based reading ability and the presence of RD, especially in terms of hemispheric dominance. Previous research has shown atypical task-based left-lateralization for this frequency range in children with RD, and intrinsic left-lateralization in typical adults. Contrary to our expectations, in our results we observed both a lack of effect of phonemic decoding ability on low-γ power/lateralization during rest, as well as possible left-lateralization in the low-γ band for the RD-only group, but not the TRs; further discussion of this finding’s robustness is warranted.

### RD effects & lateralization

At the group level, there was no evidence of RD status on low-γ power in either the left or right temporal lobes, nor was there an interaction between group and hemisphere. Moreover, although low-γ power in the left hemisphere was larger than in the right across both groups (Fig. 3), the effect of Hemisphere did not support a relative likelihood of the alternative hypothesis. The corresponding regression analyses did not show that phonemic decoding scores covaried with low-γ in either the left or right temporal regions. Rather, the Bayes factors generated from our analysis suggest that the observed data is reliably attributable to the null hypothesis. We conclude that at the population level, there are no clear differences in absolute low-γ power in the temporal regions as a function of phonemic decoding ability. We now discuss this result further in the context of hemispheric lateralization of low-γ rhythms.

Contrary to the hypotheses, the LI did not differ from 0 in the TRs, suggesting an absence of notable leftwards asymmetry in the typically-developing population during rest. The analysis in the RD-full group was less conclusive (BF ≈ 1), with there being no strong evidence for either the null or alternative hypotheses relative to one another. A follow-up test in the RD-only subgroup (excluding ADHD comorbidity) showed relatively stronger support for left-lateralization compared to null effects; this could be interpreted to reflect greater left-dominance of low-γ activity during resting-state in a population with pure phonemic decoding deficits compared to TRs or those with comorbid attentional deficits. However, an additional exploratory post-hoc analysis in the RD-full group suggested that there was no clear covariance between phonemic decoding skills and low-γ lateralization. Therefore, we conclude that any group-level left-lateralization that exists in the RDs as a whole is not strongly linked to individual differences in phonemic decoding skill.

A second exploratory post-hoc analysis suggested that for the TRs, higher phonemic decoding scores were predictive of reduced left-lateralization. However Bayesian statistical tests revealed that despite the observed data being more likely to occur under the alternative hypothesis than the null (P(data | H_1_) > P(data | H_0_)), the null hypothesis was still more likely given the observed data (P(H_0_ | data) > P(H_1_ | data)). As a result, we cannot conclude with a high degree of confidence that individual differences in phonemic decoding are associated with low-γ lateralization in our sample during rest. Nonetheless, there are a few interesting points to note from these results.

First, the lack of asymmetry for TRs may indicate that in typical decoders, there is only very minor left-lateralization at rest, if any at all, for hierarchical processing at phonemic timescales in the temporal lobes; as opposed to prior task-based studies suggesting left-hemisphere dominance for various frequency bands [40–43]. Nonetheless, this is consistent with the small effects reported for typical adults in temporal auditory and other cortical language regions at rest [12–13]. For Morillon et al. [13] in particular, results indicated that speech-sampling frequencies including the low-γ range (<47 Hz) may be expressed asymmetrically primarily during language stimulation and not at rest. We had hypothesized the possibility that in our sample of young children and adolescents, oscillations in the 30-45 Hz range may be more active at rest compared to adults, prior to much of the developmental synaptic pruning and cortical network refinement. At the same time, these children would have undergone some amount of pruning, especially compared to the pre-readers who did have resting-state left-dominance of low-gamma activity [20–21]. This paper’s null findings for lateralization in the controls therefore builds on prior ambiguous resting-state findings from pre-readers and adults. We support the perspective that asymmetric neural oscillations corresponding to phonemic timescales may facilitate linguistic processing primarily during exposure to speech, even at relatively young ages.

Second, the possibility of there being elevated left-dominance in the RD-only group, while contrary to our hypotheses, has been proposed in some previous research in relation to the ‘phonemic oversampling’ hypothesis of elevated gamma oscillations discussed by Lehongre et al. in [14]. Briefly, the authors hypothesize that both (1) reduced entrainment to acoustic modulations <30 Hz in left auditory cortex, and (2) enhanced entrainment to modulations >40 Hz in both left and right auditory cortex, can account for some of the observed deficits in RD. As our analyzed frequency range included 40-45 Hz (full range 30-45 Hz), our results are not necessarily inconsistent with (2). Lehongre et al. also report reduced left-dominance for entrained - not resting-state - oscillations in the 25-35 Hz range for RD, but do not address the 35-40 Hz range. A similarly unexpected result was reported in [44], whose authors found that increased cortical synchrony to 20 Hz modulations at ages 5-7 was negatively associated with word/pseudoword reading scores at 2-year follow-ups. There have been multiple other studies which report reduced power in RD within the phonemic frequency range, which is generally defined within the range of 20-50 Hz (see Introduction). We therefore acknowledge that a limitation of both our study and other frequency-based analyses of RD is a lack of consistency (and by extension result generalizability) of the frequency ranges ascribed to cognitive processing categories, particularly phonemic sampling. Regardless, sampling at phonemic rates in non-optimal contexts (for instance, in the absence of speech stimulation, as in this study) may be detrimental to linguistic processing if such ‘oversampling’ leads to the encoding of environmental noise [45]. Those with RD are reported to be more susceptible to the effects of noise, with many having speech-in-noise perception deficits [46–47]. Regardless, both the reported statistical results and qualitative examination of the lateralization plots (Fig. 6) make clear that intrinsic low-γ power and lateralization are endophenotypes with large individual variability that do not reliably distinguish RDs and TRs at the population level, and we do not want to overstate the degree of effect observed in this study.

Another equally important explanation to consider when evaluating these results critically is participant sampling, in particular sample sizes. An advantage of Bayesian statistical analysis is that it allows one to examine how accumulated evidence (increasing the number of data points sampled) changes the estimated probabilities of the various hypotheses. The supplementary material demonstrates the effects of such accumulated evidence in our subgroup laterality analyses (S2 Fig). It shows that for equal sample sizes of N = ∼50 in both the TR and RD-full subgroups, the accumulated evidence may have implied that the TRs demonstrate anecdotal left-lateralization, while the RD-full group does not. We can therefore appropriately modify our degree of confidence in the reported results regarding group differences by acknowledging this ambiguity.

This paper also contributes to an ongoing discussion on sampling strategies in psychology and neuroscience, especially in relation to the replicability crisis [48]. It is common to have a total N < 50, while at the same time statistical power often goes unacknowledged or unreported. Analyzing large and openly-available datasets may help with this issue somewhat, although it restricts the ability of researchers to analyze functional activity from their own task designs. Combined with the point on generalizability mentioned in the previous paragraph, we also recommend that future studies which aim to replicate or contribute to the neural oscillation literature in RD analyze multiple frequency ranges for each level of processing - for instance, if analyzing phonemic sampling, replicate each analysis for 25-35 Hz, 30-40 Hz, 30-45 Hz, 35-45 Hz, etc. This will allow researchers to investigate whether subtle variation in frequency ranges may produce different results, while removing variance due to differences in data collection and preprocessing procedures.

### ADHD comorbidity

The effects of ADHD comorbidity in our sample are worth some consideration given its high rate of occurrence in the RD population. The extent to which attentional deficits contribute to one of potentially many multifactorial pathways for the onset of reading deficits is still under debate [6]. Our results suggest a potential dissociation between the endophenotypes of ‘pure’ RD and RD-ADHD, in that left-lateralization of low-γ rhythms for the RDs was only present in the subsample with phonemic decoding deficits and no comorbid attentional deficits. It is possible that in our sample, some proportion of the RD-ADHD participants had secondary rather than primary reading deficits, and may struggle with reading due to ‘undertreated’ ADHD [6]; for the original description of the ‘phenocopy hypothesis’, see [49]. In short, impaired selective attention may negatively impact acquisition of reading skills without a characteristic ‘dyslexic endophenotype’ of atypical left-hemisphere entrainment; nonetheless, the data presented is insufficient to conclude with high confidence that this is the case for our sample.

Overall, the unique effects of attentional deficits and their relationship to reading ability are an active area of research which generate much controversy. The Multiple Deficit Model of dyslexia suggests that more comorbid conditions can confer liability to more severe deficits in cognitive and behavioral measures [50–51]. At the same time, it allows one to deduce from the high rates of comorbidity between RD and ADHD that the two disorders may overlap in their etiologies, having some shared explanatory factors that can account for behavioral deficits in both conditions and their comorbidity [52–53]. Other groups have argued that phase-resetting, entrainment, and the creation of cortical representations for stimuli may be targets of attentional modulation [54–57]. Robust selective attention during early low-level sensory processing could underlie successful entrainment, and this may be affected in children with ADHD [58–59]. However, the most robust findings of abnormal oscillatory entrainment in ADHD suggest enhanced (rather than attenuated) delta/theta and reduced absolute beta power [59–60], which we did not examine in this study - therefore, we cannot comment further on these possible effects here. There are also reports of atypical resting-state high-gamma oscillations (61-90 Hz) in adults with ADHD compared to controls [61]. Furthermore, absolute gamma power in the range of 35-45 Hz may be associated with ADHD in children [62], although results are not conclusive (for a review of resting-state EEG studies in neuropsychiatric disorders including ADHD, see [60]).

### Sex effects & their relation to RD/ADHD prevalence

The planned analyses showed that there was a large sex effect on both temporal low-γ power and lateralization: female participants had lower gamma power in both temporal gyri and increased leftwards lateralization compared to male participants. These effects held when controlling for phonemic decoding, age, and socioeconomic status. In [63], the authors showed that males 6-13 years old have a greater occurrence of resting-state EEG microstates (defined as short time periods, ∼80-120ms, during which the global scalp potential has stable topography) associated with activity in attention networks compared to females in the same age range. A later paper which also examined resting-state EEG microstates, this time in children aged 4-8 years old, further showed that the topographies of sex-dependent microstates (in which males spent more time than females) were largely localized to attention- and cognitive control-related networks [64]. Other researchers’ analyses of sex differences revealed that both processing speed and inhibitory control mediated the sex differences often reported in RD, which may also partially explain sex differences in ADHD prevalence [65].

It is possible that the reduced temporal low-γ power in females shown here may be reflective of the difference in EEG microstates indicated by past research on sex differences within this age range. However, the full implications of this are unclear. It could be that these differences reflect a developmental difference in attentional processes in boys compared to girls, as described in [65]. We therefore must acknowledge another major limitation of this paper, which is the possibility that, for the Bayesian subgroup t-tests, a higher relative proportion of boys in the RD-ADHD group, and higher relative proportion of girls in the RD-only group, could have affected the result observed. In our sample, the ratio of boys to girls in the RD-only group was close to equivalent (1.1:1); for the RD-ADHD sample, the proportion of boys was higher (1.69:1). Future research should better control for the rates of comorbidity in both males and females while maintaining statistical power, in order to determine whether the lateralization observed in the RD-only group, but not RD-ADHDs, is attributable in part to sex-based differences. The etiological pathway for these sex differences may overlap with those of attentional networks that are associated with ADHD itself, which could also partially explain the higher rate of ADHD in boys. If this is the case, then maintaining an equal ratio of boys and girls in both the RD-only and RD-ADHD groups could reduce or erase the result observed here. On the other hand, if these sex differences in attention are largely dissociated from the etiological pathway for ADHD one might expect controlling the male:female ratio not to have any effect on the result produced.

## Conclusions

The results of our analysis suggest that RD status and poor phonemic decoding do not reliably predict properties of resting-state low-γ rhythms. Results of note only emerged in the post-hoc tests, where RDs with pure decoding deficits and no comorbid attentional disorders showed a lateralization index >0 (left-dominant) for intrinsic low-γ oscillations. Similarly, an exploratory analysis in TRs revealed that greater phonemic decoding ability was associated with reduced left-laterality. Effect sizes were small, casting some doubt on the robustness of the results.

## Supporting information

**S1 Fig. Robustness checks based on prior specification for the subgroup Bayesian one-sample t-tests.**

Plots for each group t-test show how sensitive the Bayes factor is to changes in the initial Cauchy prior width (X-axis). Y-axis indicates the value of the Bayes factor with the given prior; the ordinal scale on the right (‘Evidence’) is a colloquial interpretation for the corresponding y-value. Reported results in the main text are derived from the default user prior (gray dot). (a) = TRs; (b) = RD-full; (c) = RD-only subgroup.

**S2 Fig. Sequential analysis (accumulated evidence) of the subgroup Bayesian one-sample t-tests.**

These tests indicate how the quantified degree of evidence changes as additional ‘samples’ are added. X-axis (*n*) indicates sample size. Y-axis reflects the value of Bayes factors at the given sample size; the ordinal scale on the right (‘Evidence’) is a colloquial interpretation for the corresponding y-value. Line style indicates the value of the Cauchy prior. (a) = TRs; (b) = RD-full; (c) = RD-only subgroup.

## Notes

### Competing Interest Statement

The authors have declared no competing interest.

## References

1. Peterson RL, Pennington BF. Developmental dyslexia. Annu Rev Clin Psychol. 2015;11:283–307. doi: 10.1146/annurev-clinpsy-032814-112842. Epub 2015 Jan 14. PMID: 25594880.

2. Wagner RK, Zirps FA, Edwards AA, Wood SG, Joyner RE, Becker BJ, et al. The Prevalence of Dyslexia: A New Approach to Its Estimation. J Learn Disabil. 2020 Sep/Oct;53(5):354–365. doi: 10.1177/0022219420920377. Epub 2020 May 26. PMID: 32452713; PMCID: PMC8183124.

3. Rutter M, Yule W. The concept of specific reading retardation. J Child Psychol Psychiatry. 1975 Jul;16(3):181–97. doi: 10.1111/j.1469-7610.1975.tb01269.x. PMID: 1158987.

4. Demb JB, Boynton GM, Heeger DJ. Functional magnetic resonance imaging of early visual pathways in dyslexia. J Neurosci. 1998 Sep 1;18(17):6939–51. doi: 10.1523/JNEUROSCI.18-17-06939.1998. PMID: 9712663; PMCID: PMC6792964.

5. Gayán J, Willcutt EG, Fisher SE, Francks C, Cardon LR, Olson RK, et al. Bivariate linkage scan for reading disability and attention-deficit/hyperactivity disorder localizes pleiotropic loci. J Child Psychol Psychiatry. 2005 Oct;46(10):1045–56. doi: 10.1111/j.1469-7610.2005.01447.x. PMID: 16178928.

6. Hendren RL, Haft SL, Black JM, White NC, Hoeft F. Recognizing Psychiatric Comorbidity With Reading Disorders. Front Psychiatry. 2018 Mar 27;9:101. doi: 10.3389/fpsyt.2018.00101. PMID: 29636707; PMCID: PMC5880915.

7. Langer N, Benjamin C, Becker BLC, Gaab N. Comorbidity of reading disabilities and ADHD: Structural and functional brain characteristics. Hum Brain Mapp. 2019 Jun 15;40(9):2677–2698. doi: 10.1002/hbm.24552. Epub 2019 Feb 19. PMID: 30784139; PMCID: PMC6508987.

8. Dandache S, Wouters J, Ghesquière P. Development of reading and phonological skills of children at family risk for dyslexia: a longitudinal analysis from kindergarten to sixth grade. Dyslexia. 2014 Nov;20(4):305–29. doi: 10.1002/dys.1482. Epub 2014 Sep 25. PMID: 25257672.

9. Melby-Lervåg M, Lyster SA, Hulme C. Phonological skills and their role in learning to read: a meta-analytic review. Psychol Bull. 2012 Mar;138(2):322–52. doi: 10.1037/a0026744. Epub 2012 Jan 16. PMID: 22250824.

10. Giraud AL, Poeppel D. Cortical oscillations and speech processing: emerging computational principles and operations. Nat Neurosci. 2012 Mar 18;15(4):511–7. doi: 10.1038/nn.3063. PMID: 22426255; PMCID: PMC4461038.

11. Han J, Mody M, Ahlfors SP. Gamma phase locking modulated by phonological contrast during auditory comprehension in reading disability. Neuroreport. 2012 Oct 3;23(14):851–6. doi: 10.1097/WNR.0b013e32835818e1. PMID: 22889887; PMCID: PMC4043393.

12. Giraud AL, Kleinschmidt A, Poeppel D, Lund TE, Frackowiak RS, Laufs H. Endogenous cortical rhythms determine cerebral specialization for speech perception and production. Neuron. 2007 Dec 20;56(6):1127–34. doi: 10.1016/j.neuron.2007.09.038. PMID: 18093532.

13. Morillon B, Lehongre K, Frackowiak RS, Ducorps A, Kleinschmidt A, Poeppel D, et al. Neurophysiological origin of human brain asymmetry for speech and language. Proc Natl Acad Sci U S A. 2010 Oct 26;107(43):18688–93. doi: 10.1073/pnas.1007189107. Epub 2010 Oct 18. PMID: 20956297; PMCID: PMC2972980.

14. Lehongre K, Ramus F, Villiermet N, Schwartz D, Giraud AL. Altered low-γ sampling in auditory cortex accounts for the three main facets of dyslexia. Neuron. 2011 Dec 22;72(6):1080–90. doi: 10.1016/j.neuron.2011.11.002. PMID: 22196341.

15. Lehongre K, Morillon B, Giraud AL, Ramus F. Impaired auditory sampling in dyslexia: further evidence from combined fMRI and EEG. Front Hum Neurosci. 2013 Aug 9;7:454. doi: 10.3389/fnhum.2013.00454. PMID: 23950742; PMCID: PMC3738857.

16. Cutini S, Szűcs D, Mead N, Huss M, Goswami U. Atypical right hemisphere response to slow temporal modulations in children with developmental dyslexia. Neuroimage. 2016 Dec;143:40–49. doi: 10.1016/j.neuroimage.2016.08.012. Epub 2016 Aug 9. PMID: 27520749; PMCID: PMC5139981.

17. Kershner JR. Neuroscience and education: Cerebral lateralization of networks and oscillations in dyslexia. Laterality. 2020 Jan;25(1):109–125. doi: 10.1080/1357650X.2019.1606820. Epub 2019 Apr 15. PMID: 30987535.

18. Thompson EC, Woodruff Carr K, White-Schwoch T, Tierney A, Nicol T, Kraus N. Hemispheric Asymmetry of Endogenous Neural Oscillations in Young Children: Implications for Hearing Speech In Noise. Sci Rep. 2016 Jan 25;6:19737. doi: 10.1038/srep19737. PMID: 26804355; PMCID: PMC4726126.

19. Cho RY, Walker CP, Polizzotto NR, Wozny TA, Fissell C, Chen CM, et al. Development of sensory gamma oscillations and cross-frequency coupling from childhood to early adulthood. Cereb Cortex. 2015 Jun;25(6):1509–18. doi: 10.1093/cercor/bht341. Epub 2013 Dec 10. PMID: 24334917; PMCID: PMC4428298.

20. Benasich AA, Gou Z, Choudhury N, Harris KD. Early cognitive and language skills are linked to resting frontal gamma power across the first 3 years. Behav Brain Res. 2008 Dec 22;195(2):215–22. doi: 10.1016/j.bbr.2008.08.049. Epub 2008 Sep 11. PMID: 18831992; PMCID: PMC2610686.

21. Gou Z, Choudhury N, Benasich AA. Resting frontal gamma power at 16, 24 and 36 months predicts individual differences in language and cognition at 4 and 5 years. Behav Brain Res. 2011 Jul 7;220(2):263–70. doi: 10.1016/j.bbr.2011.01.048. Epub 2011 Feb 3. PMID: 21295619; PMCID: PMC3107993.

22. Babiloni C, Stella G, Buffo P, Vecchio F, Onorati P, Muratori C, et al. Cortical sources of resting state EEG rhythms are abnormal in dyslexic children. Clin Neurophysiol. 2012 Dec;123(12):2384–91. doi: 10.1016/j.clinph.2012.05.002. Epub 2012 Jun 1. PMID: 22658819.

23. Duffy FH, Denckla MB, Bartels PH, Sandini G. Dyslexia: regional differences in brain electrical activity by topographic mapping. Ann Neurol. 1980 May;7(5):412–20. doi: 10.1002/ana.410070505. PMID: 7396420.

24. Leonard CM, Eckert MA. Asymmetry and dyslexia. Dev Neuropsychol. 2008;33(6):663–81. doi: 10.1080/87565640802418597. PMID: 19005910; PMCID: PMC2586924.

25. Alexander LM, Escalera J, Ai L, Andreotti C, Febre K, Mangone A, et al. An open resource for transdiagnostic research in pediatric mental health and learning disorders. Sci Data. 2017 Dec 19;4:170181. doi: 10.1038/sdata.2017.181. PMID: 29257126; PMCID: PMC5735921.

26. Snowling MJ, Melby-Lervåg M. Oral language deficits in familial dyslexia: A meta-analysis and review. Psychol Bull. 2016 May;142(5):498–545. doi: 10.1037/bul0000037. Epub 2016 Jan 4. PMID: 26727308; PMCID: PMC4824243.

27. Torgesen JK, Wagner RK, Rashotte CA. Test of word reading efficiency (2nd ed.). 2012. Austin, TX: Pro-Ed.

28. Cowan N, Hogan TP, Alt M, Green S, Cabbage KL, Brinkley S, et al. Short-term Memory in Childhood Dyslexia: Deficient Serial Order in Multiple Modalities. Dyslexia. 2017 Aug;23(3):209–233. doi: 10.1002/dys.1557. Epub 2017 May 12. PMID: 28497530; PMCID: PMC5540735.

29. Kubota EC, Joo SJ, Huber E, Yeatman JD. Word selectivity in high-level visual cortex and reading skill. Dev Cogn Neurosci. 2019 Apr;36:100593. doi: 10.1016/j.dcn.2018.09.003. Epub 2018 Sep 29. PMID: 30318344; PMCID: PMC6969272.

30. Nugiel T, Roe MA, Taylor WP, Cirino PT, Vaughn SR, Fletcher JM, et al. Brain activity in struggling readers before intervention relates to future reading gains. Cortex. 2019 Feb;111:286–302. doi: 10.1016/j.cortex.2018.11.009. Epub 2018 Nov 16. PMID: 30557815; PMCID: PMC6420828.

31. Willcutt EG, McGrath LM, Pennington BF, Keenan JM, DeFries JC, Olson RK, et al. Understanding Comorbidity Between Specific Learning Disabilities. New Dir Child Adolesc Dev. 2019 May;2019(165):91–109. doi: 10.1002/cad.20291. Epub 2019 May 9. PMID: 31070302; PMCID: PMC6686661.

32. Groenwold RH, White IR, Donders AR, Carpenter JR, Altman DG, Moons KG. Missing covariate data in clinical research: when and when not to use the missing-indicator method for analysis. CMAJ. 2012 Aug 7;184(11):1265–9. doi: 10.1503/cmaj.110977. Epub 2012 Feb 27. PMID: 22371511; PMCID: PMC3414599.

33. Noble KG, Wolmetz ME, Ochs LG, Farah MJ, McCandliss BD. Brain-behavior relationships in reading acquisition are modulated by socioeconomic factors. Dev Sci. 2006 Nov;9(6):642–54. doi: 10.1111/j.1467-7687.2006.00542.x. PMID: 17059461.

34. Duncan GJ, Magnuson K. Socioeconomic status and cognitive functioning: moving from correlation to causation. Wiley Interdiscip Rev Cogn Sci. 2012 May;3(3):377–386. doi: 10.1002/wcs.1176. Epub 2012 Apr 2. PMID: 26301469.

35. Pedroni A, Bahreini A, Langer N. Automagic: Standardized preprocessing of big EEG data. Neuroimage. 2019 Oct 15;200:460–473. doi: 10.1016/j.neuroimage.2019.06.046. Epub 2019 Jun 21. PMID: 31233907.

36. Bigdely-Shamlo N, Mullen T, Kothe C, Su KM, Robbins KA. The PREP pipeline: standardized preprocessing for large-scale EEG analysis. Front Neuroinform. 2015 Jun 18;9:16. doi: 10.3389/fninf.2015.00016. PMID: 26150785; PMCID: PMC4471356.

37. Winkler I, Debener S, Müller KR, Tangermann M. On the influence of high-pass filtering on ICA-based artifact reduction in EEG-ERP. Annu Int Conf IEEE Eng Med Biol Soc. 2015;2015:4101–5. doi: 10.1109/EMBC.2015.7319296. PMID: 26737196.

38. Pion-Tonachini L, Kreutz-Delgado K, Makeig S. ICLabel: An automated electroencephalographic independent component classifier, dataset, and website. Neuroimage. 2019 Sep;198:181–197. doi: 10.1016/j.neuroimage.2019.05.026. Epub 2019 May 16. PMID: 31103785; PMCID: PMC6592775.

39. Gramfort A, Luessi M, Larson E, Engemann DA, Strohmeier D, Brodbeck C, et al. MEG and EEG data analysis with MNE-Python. Front Neurosci. 2013 Dec 26;7:267. doi: 10.3389/fnins.2013.00267. PMID: 24431986; PMCID: PMC3872725.

40. Spironelli C, Penolazzi B, Vio C, Angrilli A. Inverted EEG theta lateralization in dyslexic children during phonological processing. Neuropsychologia. 2006;44(14):2814–21. doi: 10.1016/j.neuropsychologia.2006.06.009. Epub 2006 Jul 31. PMID: 16876830.

41. Spironelli C, Angrilli A. Language lateralization in phonological, semantic and orthographic tasks: a slow evoked potential study. Behav Brain Res. 2006 Dec 15;175(2):296–304. doi: 10.1016/j.bbr.2006.08.038. Epub 2006 Oct 12. PMID: 17045661.

42. Vandermosten M, Poelmans H, Sunaert S, Ghesquière P, Wouters J. White matter lateralization and interhemispheric coherence to auditory modulations in normal reading and dyslexic adults. Neuropsychologia. 2013 Sep;51(11):2087–99. doi: 10.1016/j.neuropsychologia.2013.07.008. Epub 2013 Jul 18. PMID: 23872049.

43. Nárai Á, Nemecz Z, Vidnyánszky Z, Weiss B. Lateralization of orthographic processing in fixed-gaze and natural reading conditions. Cortex. 2022 Dec;157:99–116. doi: 10.1016/j.cortex.2022.07.017. Epub 2022 Sep 29. PMID: 36279756.

44. De Vos A, Vanvooren S, Vanderauwera J, Ghesquière P, Wouters J. A longitudinal study investigating neural processing of speech envelope modulation rates in children with (a family risk for) dyslexia. Cortex. 2017 Aug;93:206–219. doi: 10.1016/j.cortex.2017.05.007. Epub 2017 May 25. PMID: 28686908.

45. Sperling AJ, Lu ZL, Manis FR, Seidenberg MS. Deficits in perceptual noise exclusion in developmental dyslexia. Nat Neurosci. 2005 Jul;8(7):862–3. doi: 10.1038/nn1474. PMID: 15924138.

46. Ziegler JC, Pech-Georgel C, George F, Lorenzi C. Speech-perception-in-noise deficits in dyslexia. Dev Sci. 2009 Sep;12(5):732–45. doi: 10.1111/j.1467-7687.2009.00817.x. PMID: 19702766.

47. Dole M, Hoen M, Meunier F. Speech-in-noise perception deficit in adults with dyslexia: effects of background type and listening configuration. Neuropsychologia. 2012 Jun;50(7):1543–52. doi: 10.1016/j.neuropsychologia.2012.03.007. Epub 2012 Mar 15. PMID: 22445915.

48. Tackett JL, Brandes CM, King KM, Markon KE. Psychology’s Replication Crisis and Clinical Psychological Science. Annu Rev Clin Psychol. 2019 May 7;15:579–604. doi: 10.1146/annurev-clinpsy-050718-095710. Epub 2019 Jan 23. PMID: 30673512.

49. Pennington BF, Groisser D, Welsh MC. Contrasting cognitive deficits in attention deficit hyperactivity disorder versus reading disability. Dev Psychol. 1993;29(3):511–523. doi: 10.1037/0012-1649.29.3.511.

50. Pennington BF. From single to multiple deficit models of developmental disorders. Cognition. 2006 Sep;101(2):385–413. doi: 10.1016/j.cognition.2006.04.008. Epub 2006 Jul 17. PMID: 16844106.

51. van Bergen E, van der Leij A, de Jong PF. The intergenerational multiple deficit model and the case of dyslexia. Front Hum Neurosci. 2014 Jun 2;8:346. doi: 10.3389/fnhum.2014.00346. PMID: 24920944; PMCID: PMC4041008.

52. McGrath LM, Pennington BF, Shanahan MA, Santerre-Lemmon LE, Barnard HD, Willcutt EG, et al. A multiple deficit model of reading disability and attention-deficit/hyperactivity disorder: searching for shared cognitive deficits. J Child Psychol Psychiatry. 2011 May;52(5):547–57. doi: 10.1111/j.1469-7610.2010.02346.x. Epub 2010 Dec 3. PMID: 21126246; PMCID: PMC3079018.

53. Kibby MY, Newsham G, Imre Z, Schlak JE. Is executive dysfunction a potential contributor to the comorbidity between basic reading disability and attention-deficit/hyperactivity disorder? Child Neuropsychol. 2021 Oct;27(7):888–910. doi: 10.1080/09297049.2021.1908532. Epub 2021 Apr 13. PMID: 33849390; PMCID: PMC8485831.

54. Lakatos P, Karmos G, Mehta AD, Ulbert I, Schroeder CE. Entrainment of neuronal oscillations as a mechanism of attentional selection. Science. 2008 Apr 4;320(5872):110-3. doi: 10.1126/science.1154735. PMID: 18388295.

55. Lakatos P, O’Connell MN, Barczak A, Mills A, Javitt DC, Schroeder CE. The leading sense: supramodal control of neurophysiological context by attention. Neuron. 2009 Nov 12;64(3):419–30. doi: 10.1016/j.neuron.2009.10.014. PMID: 19914189; PMCID: PMC2909660.

56. Lakatos P, Musacchia G, O’Connel MN, Falchier AY, Javitt DC, Schroeder CE. The spectrotemporal filter mechanism of auditory selective attention. Neuron. 2013 Feb 20;77(4):750–61. doi: 10.1016/j.neuron.2012.11.034. PMID: 23439126; PMCID: PMC3583016.

57. Zion Golumbic EM, Ding N, Bickel S, Lakatos P, Schevon CA, McKhann GM, et al. Mechanisms underlying selective neuronal tracking of attended speech at a “cocktail party”. Neuron. 2013 Mar 6;77(5):980–91. doi: 10.1016/j.neuron.2012.12.037. PMID: 23473326; PMCID: PMC3891478.

58. Ray S, Niebur E, Hsiao SS, Sinai A, Crone NE. High-frequency gamma activity (80-150Hz) is increased in human cortex during selective attention. Clin Neurophysiol. 2008 Jan;119(1):116–33. doi: 10.1016/j.clinph.2007.09.136. Epub 2007 Nov 26. PMID: 18037343; PMCID: PMC2444052.

59. Calderone DJ, Lakatos P, Butler PD, Castellanos FX. Entrainment of neural oscillations as a modifiable substrate of attention. Trends Cogn Sci. 2014 Jun;18(6):300–9. doi: 10.1016/j.tics.2014.02.005. Epub 2014 Mar 12. PMID: 24630166; PMCID: PMC4037370.

60. Newson JJ, Thiagarajan TC. EEG Frequency Bands in Psychiatric Disorders: A Review of Resting State Studies. Front Hum Neurosci. 2019 Jan 9;12:521. doi: 10.3389/fnhum.2018.00521. PMID: 30687041; PMCID: PMC6333694.

61. Dor-Ziderman Y, Zeev-Wolf M, Hirsch Klein E, Bar-Oz D, Nitzan U, Maoz H, et al. High-gamma oscillations as neurocorrelates of ADHD: A MEG crossover placebo-controlled study. J Psychiatr Res. 2021 May;137:186–193. doi: 10.1016/j.jpsychires.2021.02.050. Epub 2021 Mar 1. PMID: 33684643.

62. Barry RJ, Clarke AR, Hajos M, McCarthy R, Selikowitz M, Dupuy FE. Resting-state EEG gamma activity in children with attention-deficit/hyperactivity disorder. Clin Neurophysiol. 2010 Nov;121(11):1871–7. doi: 10.1016/j.clinph.2010.04.022. Epub 2010 May 18. PMID: 20483659.

63. Tomescu MI, Rihs TA, Rochas V, Hardmeier M, Britz J, Allali G, et al. From swing to cane: Sex differences of EEG resting-state temporal patterns during maturation and aging. Dev Cogn Neurosci. 2018 Jun;31:58–66. doi: 10.1016/j.dcn.2018.04.011. Epub 2018 Apr 30. PMID: 29742488; PMCID: PMC6969216.

64. Bagdasarov A, Roberts K, Bréchet L, Brunet D, Michel CM, Gaffrey MS. Spatiotemporal dynamics of EEG microstates in four-to eight-year-old children: Age- and sex-related effects. Dev Cogn Neurosci. 2022 Jul 12;57:101134. doi: 10.1016/j.dcn.2022.101134. Epub ahead of print. PMID: 35863172; PMCID: PMC9301511.

65. Arnett AB, Pennington BF, Peterson RL, Willcutt EG, DeFries JC, Olson RK. Explaining the sex difference in dyslexia. J Child Psychol Psychiatry. 2017 Jun;58(6):719–727. doi: 10.1111/jcpp.12691. Epub 2017 Feb 8. PMID: 28176347; PMCID: PMC5438271.

